# Neural activity for complex sounds in the marmoset medial prefrontal cortex

**DOI:** 10.1101/2024.04.22.590560

**Authors:** Rebekah E. Gilliland, Janahan Selvanayagam, Alessandro Zanini, Kevin D. Johnston, Stefan Everling

## Abstract

Vocalizations play an important role in the daily life of nonhuman primates and are likely precursors of human language. Recent functional imaging studies in the highly vocal common marmoset (*Callithrix jacchus*) have suggested that medial prefrontal cortex area 32 may be a part of a vocalization-processing network but the response properties of area 32 neurons to auditory stimuli remain unknown. Here, we performed electrophysiological recordings in area 32 with high-density Neuropixels probes and characterized neuronal responses to a variety of sounds including conspecific vocalizations. More than half of the neurons in area 32 responded to conspecific vocalizations and other complex auditory stimuli. These responses exhibited dynamics consisting of an initially non-selective reduction in neural activity, followed by an increase in activity that immediately conveyed sound selectivity. Our findings demonstrate that primate mPFC area 32 plays a critical role in processing species-specific and biologically relevant sounds.

## Introduction

Vocal communication plays a critical role among socially living species, facilitating the regulation of social interactions and coordination of group activities. The ability to produce and interpret a variety of context-specific vocalizations has been crucial in the evolutionary trajectory of primates, potentially laying the groundwork for the development of human language, which is now our primary means of communication ^1^.

The New World common marmoset monkey (*Callithrix jacchus*) is a highly vocal primate species, employing a sophisticated array of vocalizations for social interaction ^2,3^. These small-bodied primates exhibit complex vocal behaviors including vocal turn-taking and adaptive modulation of call characteristics in response to environmental variables such as background noise ^4,5^. This suggests well-developed cortical mechanisms for flexible control of vocalizations. Consequently, the neural basis of vocal communication in marmoset model has been the subject of intensive recent investigation ^6^.

Our own recent work investigated the network underlying the processing of vocalizations by using ultra-high field functional magnetic resonance imaging (fMRI) at 9.4T to map whole-brain activations in marmosets exposed to a variety of sounds, including conspecific vocalizations ^7,8^. In addition to auditory core, belt, and parabelt areas, we found strong activations for marmoset calls in mPFC area 32, suggesting that this region may play an important role the processing of species-specific sounds. Previous anatomical studies have demonstrated that this region has extensive anatomical connections with auditory cortices ^9,10^. These include superior temporal polar cortex, as well as belt and parabelt regions which consist mostly of high-order auditory association areas and respond to complex auditory stimuli including conspecific vocalizations ^11^. Because of their dense connections with auditory cortices, it has been proposed that mPFC areas 10/32v and 32 are auditory “hotspots” or even “auditory fields” ^10^. To date however, nothing is known about the auditory responses of single neurons in this region, the characterization of which is critical for understanding their role in neural circuits responsible for auditory processing.

Here, we addressed this gap by performing electrophysiological recordings in area 32 using ultra-high density 384-channel Neuropixels probes ^12^ in two marmoset monkeys presented with marmoset calls and other complex sounds. Our findings demonstrate that approximately half of the neurons in area 32 respond to these stimuli. The initial response to the sounds was a non-selective reduction in neural activity, followed more than 100 ms later by an increase in activity that immediately conveyed sound selectivity.

## Results

We recorded neural activity in response to eleven complex sounds in the medial prefrontal cortex (mPFC) of two adult female marmosets using high-density Neuropixels probes. Ex-vivo ultra-high field magnetic resonance imaging confirmed that the probes targeted the mPFC, with most recordings likely within areas 32 and 32v, and a minority just dorsal to area 32 in area 9. A few recordings may have extended ventrally into area 14R (Fig. 1A). We conducted 12 sessions with marmoset L and 6 sessions with marmoset D, and recorded the activity of 2,213 and 563 neurons, respectively.

**Figure 1.**
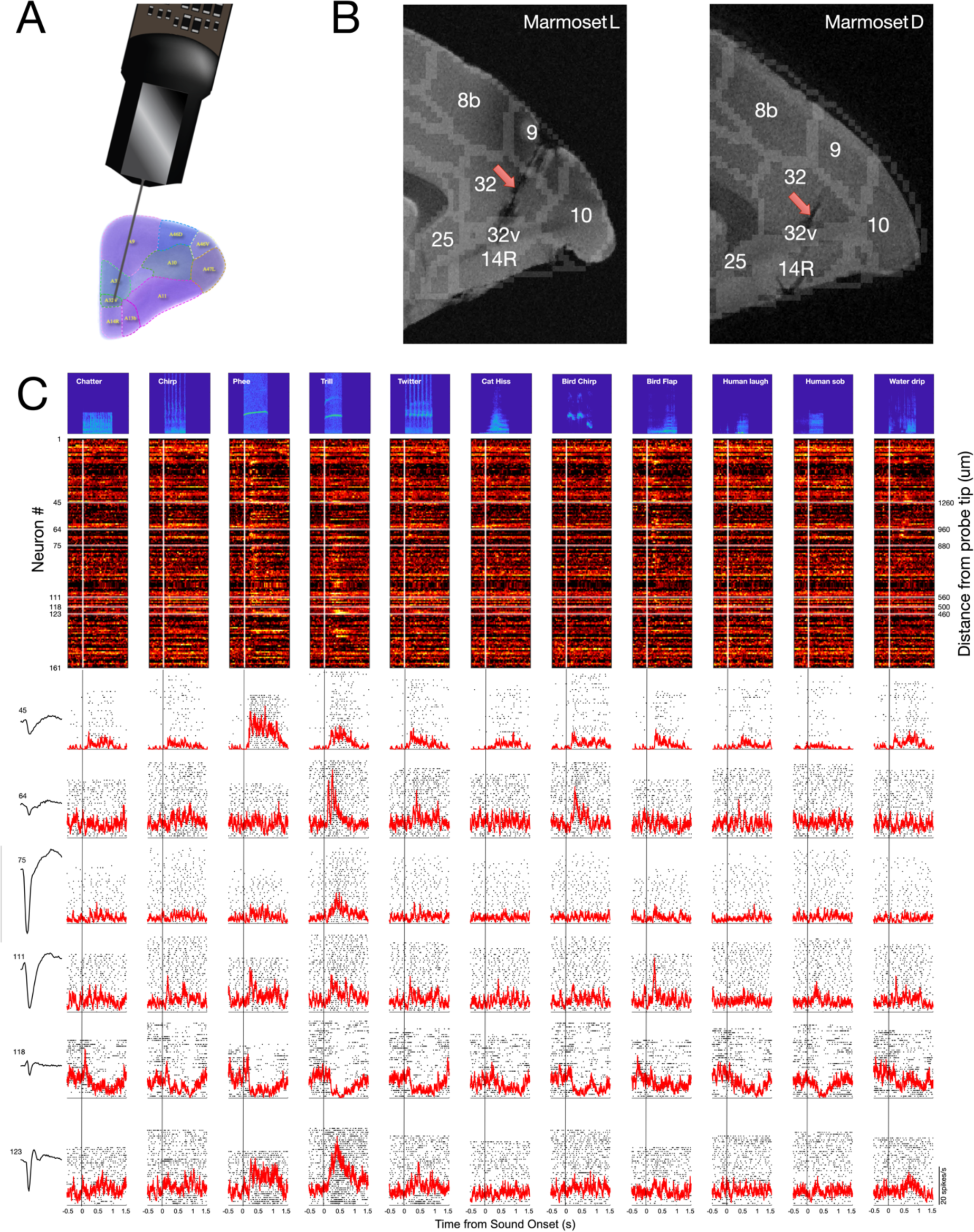
Recording approach, confirmation of recording sites, and auditory response profiles of example neurons. (A) Schematic depiction of Neuropixels probe and trajectory to area 32. (B) ex-vivo anatomical MRI images in sagittal plane obtained at 15.2T showing location of electrode tracts (red arrows) and termination within areas 32 and 32v of marmoset medial frontal cortex. (C) Representative auditory responses of single area 32 neurons to 11 sound stimuli, recorded simultaneously within a single recording session. Top panel, spectrograms of auditory stimuli presented, including marmoset vocalizations. Middle panel, auditory responses of single area 32 neurons aligned on auditory stimulus onset and sorted by depth from probe tip. Each row depicts mean discharge rate within 20 ms bins for a single area 32 neuron, colourmap indicates normalized response magnitude. Lower panel, responses of single neurons to the suite of 11 auditory stimuli. Rasters and spike density functions aligned on stimulus onset. Mean action potential waveforms for each example neuron shown at left.

Figure 1B depicts the normalized responses to the 11 sounds from 161 neurons recorded simultaneously in a single session in marmoset L. In this session, many neurons responded robustly to phee and trill calls. The activity of six example neurons is shown in Figure 1C. These neurons were recorded within an 800 μm range (see Fig. 1B for their recording depth relative to the probe tip). The neuron in the top row showed a preference for phee calls, whereas the neurons in the second and third rows exhibited higher activity for the trill call. While the neuron in the fourth row also responded to both phee and trill calls, the greatest response was to the sound of beating bird wings. We also observed neurons that decreased their activity in response to the sounds (see Fig. 1B). An example of this inhibitory response is plotted in the fifth row. The neuron in the last row responded to both trill and phee calls. The first four neurons exhibited broad action potentials, and the other two had narrow action potentials (Fig. 1C, left), patterns typically associated with putative pyramidal neurons and interneurons, respectively.

We found that some neurons had extremely low discharge rates. For all subsequent analyses, we included only neurons that discharged at least one spike per second for any of the 11 sounds (see Methods). A total of 1,908 neurons (1,448 from marmoset L and 460 from marmoset D) met this criterion. To determine if neurons were responsive to the sounds, we compared the activity to all sounds against baseline activity using paired t-tests (see Methods). To evaluate selectivity for the sounds, we performed one-way ANOVAs on the baseline-corrected activity for the set of 11 sounds for each neuron. In marmoset L, 849 neurons were responsive and 446 were selective (30%). Out of 1,448 neurons, 991 (68%) were either responsive or selective. In marmoset D, we found 260 responsive and 101 selective neurons, with 291 of 460 neurons (or 63%) displaying either responsiveness or selectiveness for the sounds. Across both marmosets, 1,282 of 1,908 neurons (67%) responded to the sounds or showed selectivity. Of these 1,282 neurons, 547 (or 43%) were sound selective. This finding is consistent with our previous fMRI results which indicated strong activation for sounds in marmoset area 32.

As illustrated in Figures 1B and C, we found that some neurons increased their activity following presentation of complex sound stimuli, while the activity of others decreased. Based on their overall response to all the sounds compared to baseline activity (see Methods), we classified neurons as excited if their activity increased in response to these sounds (655/1,282, or 51%) or as inhibited if their activity was reduced following sound presentation (627/1,282, or 49%). Figure 2A shows the normalized activity of all excited neurons for the sound stimulus that evoked the neurons’ maximal response. Figure 2B shows the normalized activity of all inhibited neurons for the sound stimulus that drove the strongest activity suppression. Neurons were sorted based on the onset times of their responses to the sounds (see Methods). To explore whether excitatory and inhibitory responses mapped onto specific neuron types, we classified neurons as narrow (NS) (<400 μs trough-to-peak action potential duration) or broad-spiking (BS) (≥400 ms trough-to-peak action potential duration) and investigated the proportions of neurons with excitatory and inhibitory responses within each group. Figure 2B shows that both excited and inhibited neuron groups contained NS and BS neurons, with approximately 20% being NS neurons (excited: 130 NS and 525 BS neurons; inhibited: 115 NS and 512 BS). This suggests that the categorization of neurons into those that increase and those that decrease their activity in response to sounds is not directly related to the physical characteristics of their action potentials.

**Figure 2.**
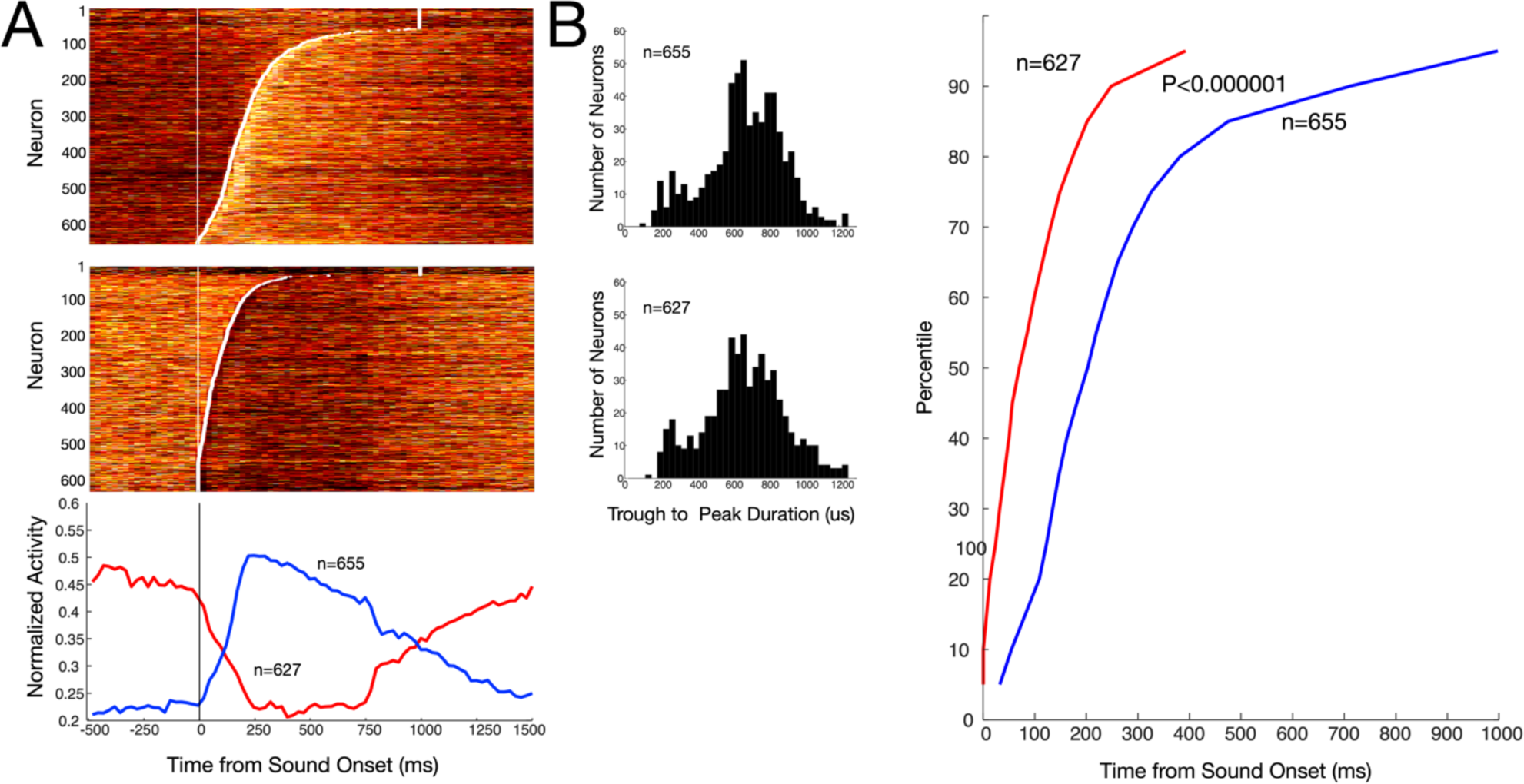
Timing and magnitude of neurons with excited and inhibited responses to auditory stimuli in area 32 neurons. (A) Discharge rates of neurons with excited (top panel) and inhibited (middle panel) responses to auditory stimuli. Each row depicts activity of a single neuron, heatmap indicates normalized mean discharge rate in 25ms bins. Neurons are sorted in order of onset times of auditory responses, depicted by white dots. Bottom panel, mean spike density functions depicting discharge rates for the subpopulations of area 32 neurons with excited (blue line) and inhibited (red line) responses. **B)** Histograms depicting trough-to-peak waveform durations for neurons with excited (top panel) and inhibited (bottom panel) responses. Proportions of broad and narrow spiking neurons did not differ between response types. (C) Cumulative onset times of auditory responses for neurons with inhibitory (red line) and excitatory (blue line) responses.

To investigate the temporal dynamics of excitatory and inhibitory responses we determined and compared their onset times. Figure 2C plots the cumulative onset of neural responses for neurons with excitatory (blue) and inhibitory (red) responses. The median onset time for excited neurons was 203 ms, whereas for inhibited neurons, it was considerably shorter at 70 ms. The onset times of these two groups were significantly different (Kruskal-Wallis test, p<0.000001), indicating that the initial response to a sound was a rapid suppression of existing neural activity, followed by a later increase in activity in a different group of neurons.

To determine whether these neurons exhibited preferences for specific sounds, we next identified the preferred sound for all selective neurons. Excited neurons predominantly preferred the flapping bird wing sound and the phee call (Figure 3A), with the weakest responses evoked by the two tested human vocalizations. Chi-square tests confirmed that these preferences were significantly non-uniform (187.6, p<0.0000001 for the sounds eliciting maximal responses, and 91.35, p<0.0000001 for the sounds eliciting minimal responses). Among neurons with inhibited responses (Figure 3B), many showed a preference (i.e., decreased their activity) for flapping bird wing sounds and also marmoset calls (Chi-square test: 37.13, p<0.0001), with the weakest responses (i.e., largest activity) towards cat hisses and bird chirps (Chi-square test: 38.9, p<0.0001). Overall, these neurons preferred the sound of flapping wings and phee and trill calls.

**Figure 3.**
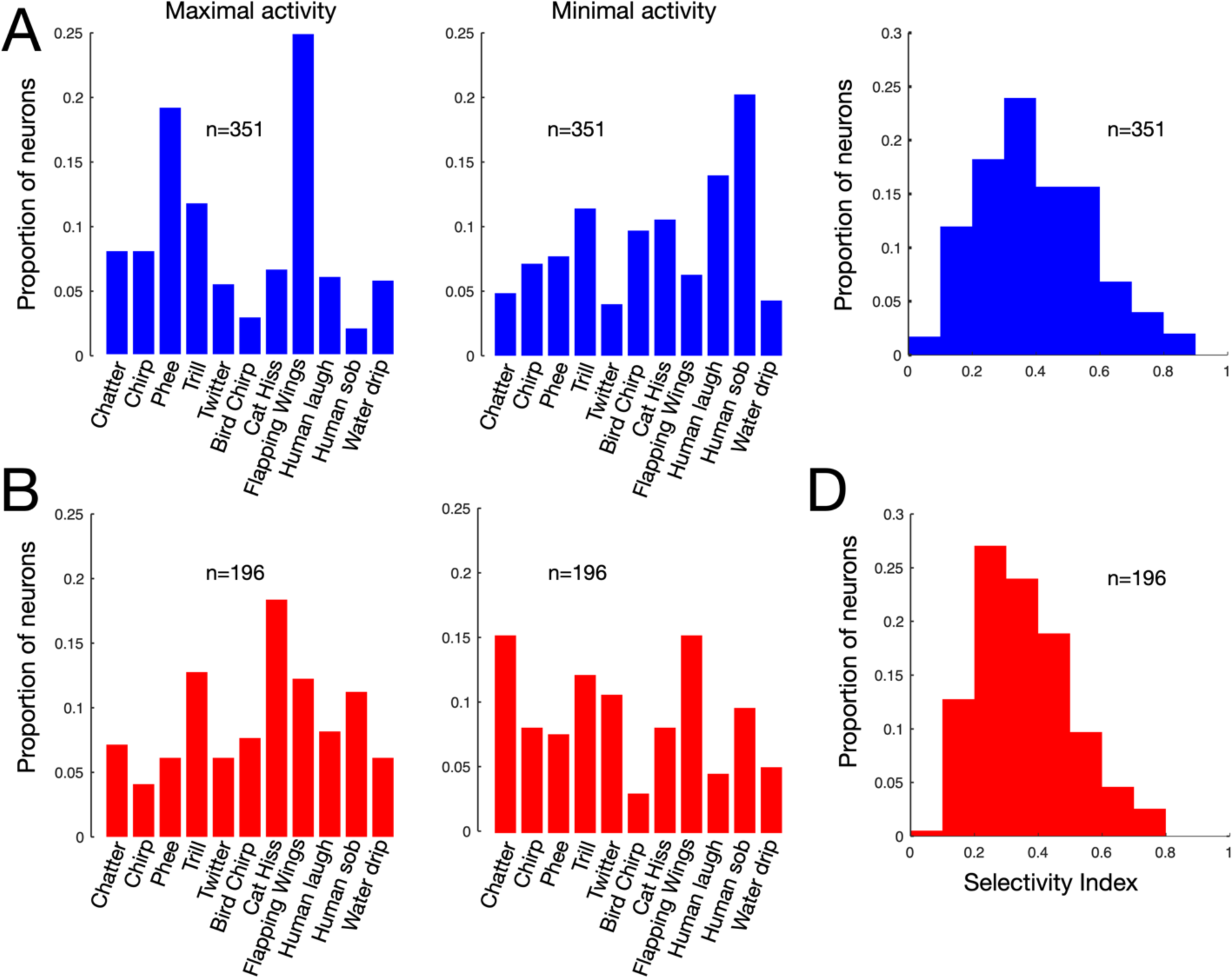
Selectivity of excited and inhibited responses of area 32 neurons to auditory stimuli. (A) Left panel, proportion of neurons with excited responses that exhibiting maximal increase in activity for each of the 11 sounds, Right panel, same for minimal activity for each sound. (B) Same as A for neurons with inhibited responses. (C) Histogram of selectivity indices computed for the suite of presented sounds for excited neurons. (D) Same as C for inhibited neurons.

To characterize the sound selectivity of these neurons, we computed a selectivity index for each neuron (see Methods). Index values close to zero indicate that a given neuron responded similarly to all 11 sounds, whereas values approaching one would indicate an exclusive response to one of the eleven sounds. We observed a range of selectivity indices for both excited (Figure 3C) and inhibited neurons (Figure 3D), with excited neurons having a mean index of 0.39 and inhibited neurons having a mean index of 0.36. Selectivity indices between the two groups differed significantly (Wilcoxon Rank Sum test, p=0.038), a finding potentially reflecting the reduced dynamic range of neurons in which the predominant response was a decrease from baseline.

For a subset of neurons, we additionally recorded their responses during presentations of scrambled versions of the original 11 sounds (see Methods). Figure 4A illustrates the responses to original and scrambled sounds for the same neuron shown the top row of Figure 1B. The neuron showed a preference for the intact phee call but also responded to the scrambled phee call, albeit less strongly. Figure 4B displays the selectivity indices of all excited neurons for intact calls plotted against the selectivity indices for the 11 scrambled calls. For the 281 excited neurons, their mean selectivity decreased from 0.41 for the intact sounds to 0.35 for the scrambled sounds (paired t-test, p<0.000001). A similar trend was observed for the 137 inhibited neurons (Figure 4C), where the mean selectivity index dropped from 0.39 for intact sounds to 0.32 for the scrambled sounds (paired t-tests, p<0.000001). These results suggest that both subsets of neurons are more adept at discriminating intact sounds than scrambled sounds.

**Figure 4.**
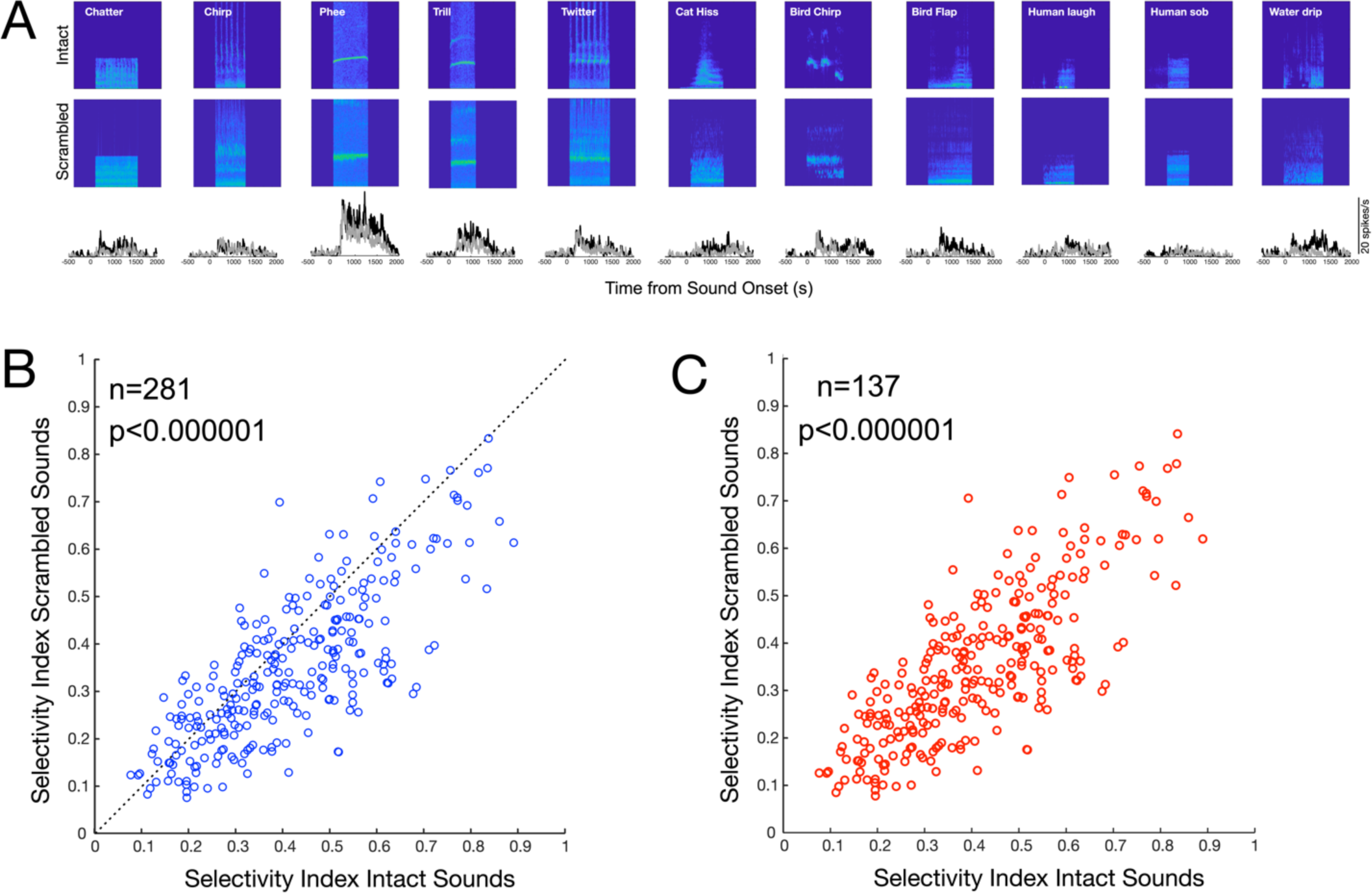
Comparison of responses of area 32 neurons to intact and scrambled sound stimuli. (A) Top two rows, spectrograms for intact and scrambled versions of the 11 auditory stimuli. Bottom row, spike density functions of responses of example neuron to intact (black lines) and scrambled (grey lines) stimuli aligned to sound onset. (B) Scatterplot of selectivity indices for scrambled and intact sounds for neurons with excited responses. (C) Same as B for neurons with inhibited responses.

**Figure 5.**
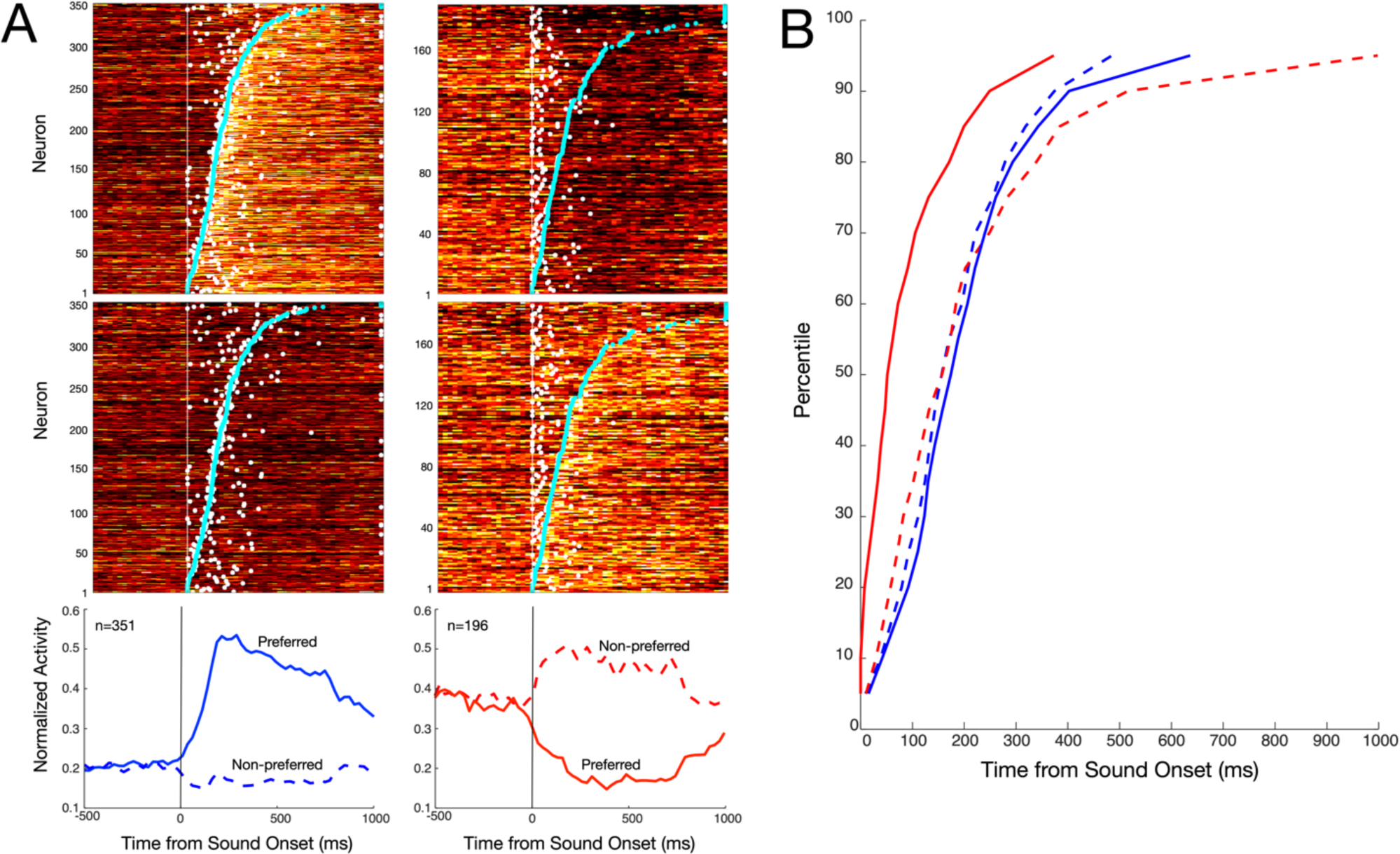
Timecourse of auditory response selectivity of neurons in area 32. (A) Top four panels, plots of mean discharge rates of single neurons aligned on auditory stimulus onset, sorted in increasing order of discrimination times (cyan dots) between their preferred and non-preferred stimuli. White dots depict onset times of auditory responses. Colourmap depicts normalized mean discharge rate computed in 25ms bins. Top left, responses of excited neurons to preferred stimulus. Middle left, responses of excited neurons to non-preferred stimulus. Top right, responses of inhibited neurons to preferred stimulus. Middle right, responses of inhibited neurons to non-preferred stimulus. Bottom panels depict mean spike density functions for preferred (solid lines) and non-preferred (dashed lines) auditory stimuli aligned on stimulus onset for excited (blue lines, bottom left) and inhibited (red lines, bottom right) neurons. (B) Cumulative plot of distribution of onset times (solid lines) and discrimination times (dashed lines) for excited (blue) and inhibited neurons (red). Discrimination times lagged onset times for inhibitory neurons only.

Finally, to examine the timecourse of response selectivity, we compared the discrimination times of excited and inhibited neurons, i.e., the time at which a neuron first exhibited significant differences in activity between its preferred and nonpreferred sound. Figure 4A shows the activity of all selective neurons aligned on sound onset and sorted by their discrimination times (cyan) of their preferred sound (top) versus their nonpreferred sound (bottom) for excited (left) and inhibited neurons (right). White dots indicate the onset times of responses - the time at which their activity first differed significantly different from baseline. Similar to all responsive neurons (Fig. 2), inhibited neurons responded to the sounds in advance of excited neurons (median 52 ms for inhibited vs. 174 ms median for excited neurons, Wilcoxon Rank Sum test, p<0.0000001). The median discrimination time (159 ms) was significantly later than the median onset time for inhibited neurons (Wilcoxon Signed Rank test test, p<0.00000001). For excited neurons, the discrimination time (155 ms) and onset time (174 ms) were not statistically different (Wilcoxon Signed Rank test p=0.12). The discrimination times between excited and inhibited neurons also did not differ significantly (Wilcoxon Rank Sum test, p=0.99), indicating that the initial response to the sounds was a non-selective reduction in neural activity in the population of neurons with inhibitory responses, followed at a latency of more than 100 ms by an increase in activity of neurons with excitatory responses that displayed immediate sound selectivity.

## Discussion

Primate medial prefrontal cortex area 32 has been posited as a prefrontal auditory field based on its extensive connections with temporal auditory cortices ^9,10,13,14^. Recent functional imaging studies in awake marmoset monkeys have supported this hypothesis by revealing activations in area 32 in response to conspecific vocalizations ^7,8^. Here, we present what we believe to be the first characterization of single neuron activity in response to complex auditory stimuli in this area and note several key findings consistent with this notion.

The observation that approximately half of the neurons in area 32 responded to complex auditory stimuli provides strong support for the consideration of area 32 as a prefrontal auditory field. In addition, the fact that roughly 40% of these auditory-responsive neurons exhibited sound selectivity is consistent with our prior fMRI findings, which showed heightened activations for vocalizations compared to non-vocal sounds or scrambled vocalizations. Taken together with the established anatomical connectivity of area 32 with cortical fields in auditory cortex ^9^, our findings here support a crucial role for mPFC area 32 in the processing of species-specific sounds. We thus propose that this area is a critical node in the network instantiating the neural processes underlying social communication in marmosets. We found additionally that marmoset area 32 neurons responded robustly to at least one non-social sound stimulus. The preferential response of neurons in this area to flapping wing sounds as well as phee and trill calls, along with the demonstrated sound selectivity, suggests a specialized processing capacity for the processing of biologically relevant sounds. Such a capability is essential for adaptive behavioral responses to both predators ^15,16^ and social cues from conspecifics ^17^. Finally, the comparison of neural responses to original and scrambled sounds further elucidates the neural basis of sound discrimination in marmosets. The observed decrease in selectivity for scrambled sounds suggests that the integrity of the sound structure may be essential for effective neural discrimination and that the marmoset’s auditory system is finely tuned to the specific features of conspecific vocalizations.

Our major finding, however, was the observed dynamics of neuronal responses following sound stimuli within area 32, in which an initially non-selective reduction in neural activity was followed by an increase in activity that conveyed sound selectivity. Retrograde tracer injections have revealed that in addition to projections from other prefrontal areas ^18^, area 32 receives significant input from two other primary sources: the basolateral (BLA) and basomedial amygdala (BMA), and higher auditory and association areas, such as the temporopolar area (TPO) and the superior temporal region (STR) ^19^. In rodents, the excitatory projection from the amygdala is known to activate local circuit inhibitory interneurons in the mPFC which then are thought to suppress neural activity within the area ^20^. This proposed pathway, wherein the BLA and BMA rapidly activate inhibitory interneurons that then suppress activity in area 32 could serve as an initial filtering mechanism, reducing ongoing neural processing and preparing the neural circuitry for more selective processing of biologically and socially relevant auditory cues. Independent lines of evidence have associated the amydala with the regulation of emotional processes such as anxiety^21^, modulation of social behaviors such as grooming ^22^, and cognitive processes including reward processing ^23^. Recently, vocalization-related activity has been identified in the marmoset BLA and BMA^24^. Based on this, an intriguing possibility is that area 32 is a site of convergence of such signals with incoming auditory stimuli, and indeed this would be consonant with the role of this area in emotional communication proposed previously ^18,25^.

The sound-selective activity we observed was present nearly immediately in the responses of area 32 neurons with excitatory responses – these neurons discriminated their preferred from non-preferred auditory stimuli within their initial stimulus-driven responses. This differs considerably from the dynamics observed in areas such as lateral PFC during selection of visual targets, in which an initially indiscriminate volley of activity evolves to discriminate target stimuli over a timecourse of tens of milliseconds ^26^. Differences in stimulus modality notwithstanding, this suggests that the excitatory sound-selective responses we observed here were driven by inputs already carrying this signal, originating most likely from higher-order temporal auditory areas which as noted previously share extensive interconnections with area 32 ^9^. These regions have been shown to be involved in integrating complex sensory information and play a crucial role in recognizing and interpreting conspecific vocalizations ^11^. The initial non-selective reduction in neural activity, followed by the aforementioned increase in activity conveying sound selectivity, suggests a hierarchical processing model wherein initial rapid filtering is followed by more detailed analysis of sound properties (Fig. 6). The interplay between inhibitory and excitatory inputs likely facilitates the discrimination and prioritization of sounds, enabling marmosets to attend and process behaviorally relevant sounds.

**Figure 6.**
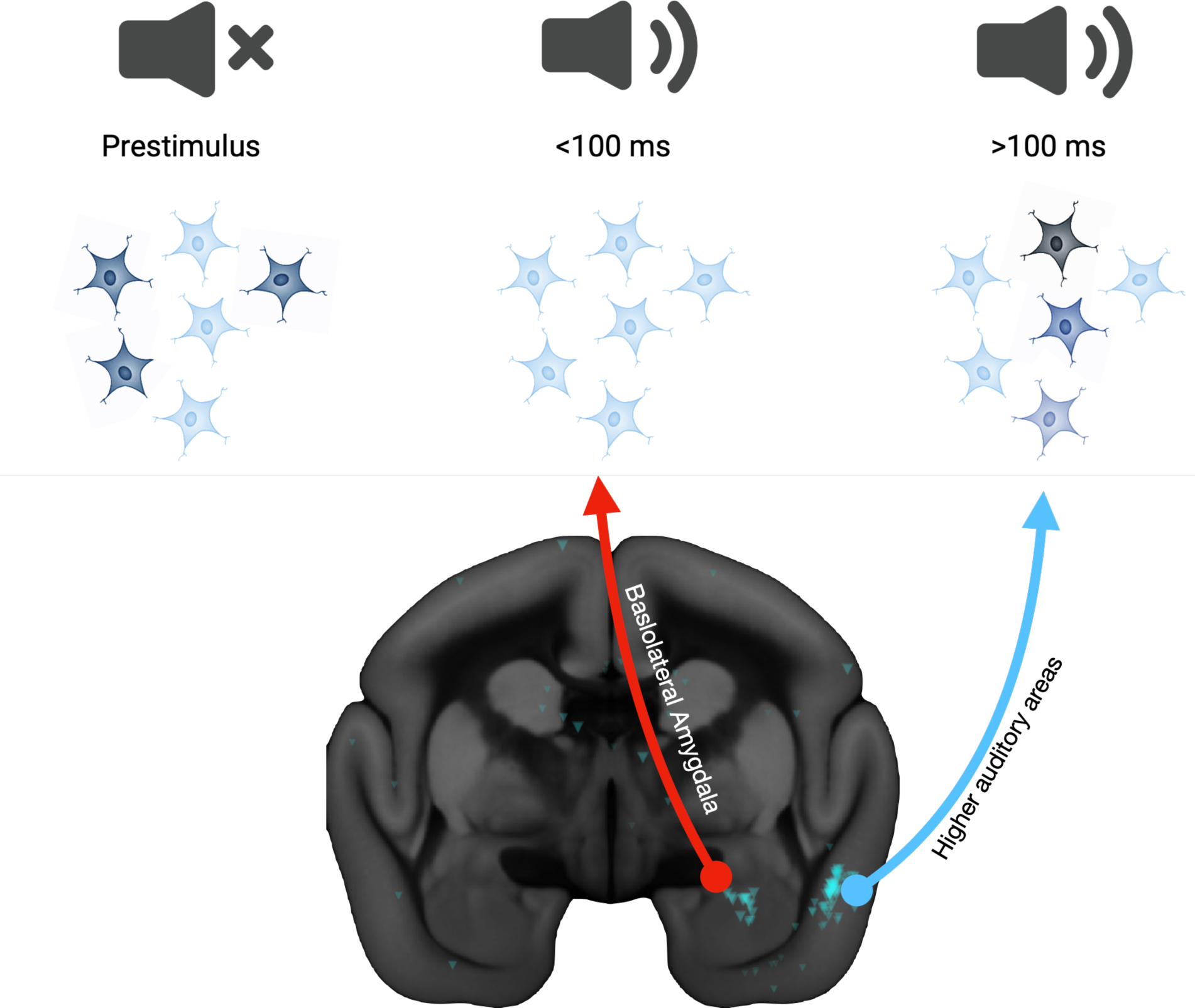
Schematic of proposed model of hierarchical processing circuit for auditory stimuli between basolateral amygdala (BLA), higher auditory areas, and area 32. Top left. Prior to onset of an auditory stimulus, many area 32 neurons exhibit spontaneous activity. Following onset of an auditory stimulus, input from amygdala neurons silences area 32 neurons (red arrow, top middle), effectively reducing noise in discharge rates. Top right, responses and selectivity of area 32 neurons are enhanced by later input from higher auditory areas (light blue, top right panel). MRI shows pre-processed retrograde tracer injection into area 32 (R01_0095) from BMCR-explorer) ^19^.

Medial prefrontal areas are interconnected with an extensive array of cortical and subcortical areas subserving cognitive, sensory, motor, and limbic functions, and it has been proposed that mPFC acts to integrate these various processes in the service of adaptive behavior^27^. Here, we found evidence supporting a role of mPFC area 32 in auditory processing of complex stimuli including conspecific vocalizations, and response dynamics consistent with known inputs from the BLA and higher-order auditory cortices. Further investigations using targeted manipulations of the BLA and connections between area 32 and higher auditory association areas will be needed to provide direct evidence for the interplay between these areas and establish the neural mechanisms underlying their interactions. Given the additional connectivity of area 32 with areas involved in vocal production ^28^, and previously proposed role in emotional vocalizations ^18,25,29,30^, it seems reasonable to suggest that a specific role of area 32 in auditory processing and communication would fit well within the generally integrative conceptual framework of mPFC function. An understanding of the mechanisms governing interactions between mPFC area 32, the BLA, and auditory cortices has the potential to significantly advance our knowledge of the neural basis of communication and its evolution, shedding light on the networks that are fundamental to social behavior in primates, including humans.

## Methods

### Subjects

Data were obtained from two female adult common marmosets (Marmoset L, age 26 months, weight 370g; Marmoset D, age 58 months, weight 390g). Prior to participation in these experiments, both animals were acclimated to restraint in a custom-designed marmoset chair ^31^. All experimental procedures were conducted in accordance with the Canadian Council on Animal Care policy on the care and use of laboratory animals and a protocol approved by the Animal Care Committee of the University of Western Ontario Council on Animal Care. The animals were additionally under the close supervision of university veterinarians throughout all experiments.

### Surgical Preparation of Animals for Electrophysiological Recordings

Marmosets underwent an aseptic surgical procedure in which a microdrill was used to open an approximately 2mm trephination allowing access to underlying cortex lateral to the sagittal sinus and above area 32 based on stereotaxic coordinates ^32^, This trephination was sealed with silicone (Kwik-Sil, WPI International, Sarasota, FLA, USA), and a combination recording chamber/head restraint was fixed to the skull using a combination of universal dental adhesive (All-Bond Universal, Curion, Richmond, BC, Canada) and UV-cured dental resin cement (Core-Flo DC, Curion, Richmond, BC, Canada). This allowed access to cortex and stabilized the head for electrophysiological recordings. Finally, the recording chamber was covered with a protective cap. Detailed descriptions of these procedures have been reported previously ^31,33^.

### Auditory Stimulus Presentation

In all sessions, marmosets were seated in a custom-designed primate chair with the head restrained^31^ inside an electrically-shielded sound attenuating chamber (Crist Instrument Co. Hagerstown MD). A spout was placed at animals’ mouths to allow delivery of a viscous liquid reward (Acacia gum powder mixed in a 50:50 ratio with water) via an infusion pump (Model NE-510, New Era Pump Systems, Inc., Farmingdale, New York, USA). Rewards were delivered to maintain alertness of the animals during each session. A CRT monitor (ViewSonic Optiquest Q115, 76 Hz non-interlaced, 1600 x 1280 resolution) was positioned in front of the animals to allow for presentation of videos depicting natural scenes during electrode advancements, and to provide background illumination during auditory stimulus presentations. In both cases, this visual stimulation was provided as an additional aid to maintain the alertness of the animals. Auditory stimulus presentations were controlled by Monkeylogic ^34^ running on an ASUS UX430U Notebook PC using the Windows 10 Operating System. Stimuli were played through a Bose Soundlink III speaker (Bose Corporation, Framingham Mass.) connected to the audio output of the Notebook PC and placed at a distance of 10cm centred in front of the animals, and 12cm below head level. For the complex stimulus task, marmosets were presented with a suite of 11 different auditory stimuli that included 5 marmoset calls (phee, trill, twitter, chirp, and chatter), bird song, flapping bird wings, cat hiss, human sigh, human laugh, and water drip. In some session, we also presented scrambled versions of these sounds. Normalization and scrambling was performed using published MATLAB code ^35^. Stimulus duration ranged from 550ms to 1000ms, with shorter stimuli reflecting naturally shorter marmoset call types, such as trill calls. Spectrograms of each normalized intact and scrambled stimulus can be seen in Figure 1. Each trial of the passive auditory task consisted of a single stimulus presentation, which was followed by a variable intertrial interval of 2000-3000ms. Stimuli were presented in random order and repeated at least 20 times within each recording session.

### Electrophysiological recordings

Electrophysical recordings were conducted using Neuropixels 1.0 NHP short probes ^12^, with the external reference and ground bridged for all recordings. All recordings were referenced to the reference contact at the tip of the electrode. Neural spikes were sampled at 30 kHz and high-pass filtered at 300 Hz. Custom Neuropixels electrode holders designed to interface with the dovetail structures on the probe base (Imec, Belgium) were fixed to Narishige Stereotaxic Micromanipulators (Narishige SM-25A, Narishige International USA Inc., Amityville, NY, USA) to advance and retract electrodes in all recordings. An IMEC headstage was used with a PXIe-8381 acquisition module. The PXIe-1082 chassis and the MXIe interface were used for data acquisition. 8-bit digital event signals emitted by Monkeylogic were recorded using the PXI-6133. Neural and auxiliary signals were synchronized by a TTL pulse emitted by Monkeylogic at target onset. All data was acquired using the SpikeGLX application (v20190413-phase3B2, Karsh, 2019).

To minimize infection risk and to improve the stability of recordings, we advanced the Neuropixels probe through the intact silicone seal in all sessions. Maintaining this seal minimized exposure of the dural surface and additionally reduced cortical pulsation. Probes were soaked in 70% ethanol for a minimum of 30 min prior to insertions. For each recording session, we followed the protocol below. We first removed the chamber cap and thoroughly rinsed the inside of the recording chamber with 0.9% sterile saline delivered via a sterile syringe with a flexible sterile catheter tip. This was soaked up with sterile gauze, and the chamber was subsequently filled with 10% iodine solution. We then scrubbed the outside of the chamber with 70% isopropyl alcohol wipes, soaked up the iodine solution with sterile gauze and repeated the saline flushing procedure. After preparing the inside of the chamber and surface of the silicone seal, we advanced the probe through silicone and dura using a stereotaxic micromanipulator. In all cases the probe was driven at an angle 10° from vertical, avoiding the large sagittal sinus and crossing of the midline. A schematic depiction of our electrode approach is presented in Fig. 1A. We continuously monitored neural activity on the probe, noting the depth at which activity was first observed at the probe tip as the cortical surface as a guide to the depth at which area 32 was reached. Once in place, we allowed the probes to settle for 30 minutes to minimize drift during the recording session. At the end of the study, we confirmed the location of recording sites in area 32 with *ex-vivo* structural MRI at 15.2T (see *Confirmation of Recording Sites with ex-vivo MRI*, below). Electrodes were allowed to settle for 30-45 minutes to minimize drift during the recording session. Animals viewed videos of naturalistic scenes during electrode advancement and the settling time to minimize restlessness. Following this, the animals were presented with the passive auditory task, the duration of which was 30-45 minutes. At the end of each session, the Neuropixels probe was carefully retracted and the integrity of the silicone seal was checked. This was replaced if necessary. We then replaced the chamber cap.

### Automated Spike Sorting with Manual Curation

Putative single unit clusters were extracted using Kilosort 2 ^36^. A brief description of this approach is as follows. A common median filter was applied across channels and a “whitening” filter was then applied to reduce correlations between channels and maximize local differences among nearby channels. Following these preprocessing steps, templates were constructed based on an initial segment of the data and adapted throughout the session with an accommodation for drift over time. Clusters were then separated and merged as necessary. Following this initial automated step, putative single unit clusters were curated manually using Phy (Rossant, 2019). Here, we visualized and merged or split clusters based on inspection of waveforms, cross-correlograms, and distributions of spike amplitudes. Following manual merging and splitting of clusters, clusters with consistent waveforms, normally distributed amplitudes, a dip in the autocorrelogram at time 0, and a consistent presence throughout the recording session were marked as single units, and all others were marked as multi-unit clusters or noise clusters as appropriate, and were discarded from further analysis.

### Ex-vivo MRI

Following data collection, marmosets were euthanized and perfused to allow for reconstruction of electrode tracts using ex-vivo structural MRI’s. Animals were deeply anesthetized with 20 mg/kg ketamine plus 0.025 mg/kg medetomidine and 5% isoflurane in 1.4–2% oxygen to reach beyond the surgical plane. This was confirmed via lack of responses to physical stimulation including toe pinching and cornea touching. They were then perfused transcardially with 0.9% sodium chloride solution, followed by 10% buffered formalin. The heads were stored in 10% buffered formalin for at least 7 days prior to MRI scanning. Ex-vivo MRI of the brains were performed on a 15.2T, 11-cm horizontal bore magnet (Bruker BioSpin Corp, Billerica, MA) and Bruker BioSpec Avance Neo console with the software package Paravision360.3.5 (Bruker BioSpin Corp, Billerica, MA). The Bruker gradient/shim coil (B-GA6S-100) had a 6-cm inner diameter, a maximum gradient strength and slew rate of 1,000 mT/m and 9,000 T/m/s, respectively, and a full set of 3rd-order shims. The radiofrequency coil was a 35-mm inner diameter, transmit/receive volume coil with quadrature detection (Bruker BioSpin Corp, Billerica, MA). The sample was placed in a 50 ml conical centrifuge tube, filled with Fomblin oil (ECL, Aurora, IL), and placed under vacuum for 30 min to remove air bubbles. High resolution (50×50×50 µm) T2-weighted images were acquired for each brain. The raw MRI images were converted to NifTI format using dcm2niix ^37^ and the MRIs were manually registered to the ultra-high-resolution ex-vivo NIH template brain ^38^, that contains the location of cytoarchitectonic boundaries of the Paxinos atlas ^3232^. These cytoarchitectonic boundaries of the atlas were overlayed on the registered T2 anatomical images of the medial prefrontal cortex to identify the location of the Neuropixels probe tracts which were readily visible in each brain (Fig. 1A).

## Data Analysis

For displaying continuous spike density waveforms, spike trains were convolved with a postsynaptic activation function with a binwidth of 1 ms. This asymmetric activation waveform is designed to mimic an excitatory postsynaptic potential.

Neurons were classified as responsive if their activity was different between a baseline window 500-0 ms prior to sound and a window 50-800 ms from sound onset, evaluated by paired Student t-tests at p<0.05. Neurons that showed a higher activity during the poststimulus window were categorized as neurons with excitatory responses and neurons that displayed a lower activity during the poststimulus window were categorized as inhibited neurons. Sound selectivity was assessed using one-way ANOVAs on the baseline-corrected neural activity for the 11 intact sounds.

As a measure of stimulus selectivity, a selectivity index was defined as the following:

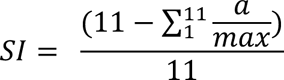

Here a= activity for current condition in the window 50-800 ms after stimulus onset; max = maximal activity To determine the onset of neural responses following sound presentation, we measured neural activity for each neuron for their preferred stimulus in by 1 ms sliding 50 ms windows. The onset time for each neuron was defined as the beginning of the first 50 ms window in which the activity became significantly greater than baseline for excited neurons and lower than baseline for inhibited neurons (evaluated by paired Student t-test at p<0.025).

To determine the discrimination time for each neuron, we compared neural activity between the preferred and nonpreferred sounds for each neuron in 50 ms sliding windows. The discrimination time was defined as the beginning of the first 50 ms window when the activity became significantly different between preferred and nonpreferred sounds (evaluated by non-paired Student t-test at p<0.025).

## Acknowledgments

We wish to thank Cheryl Vander Tuin, Whitney Froese, Miranda Bellyou, and Hannah Pettypiece for animal preparation and care and Dr. Joseph Gati for scanning assistance. Support was provided by the Canadian Institutes of Health Research (FRN 148356, S.E.) the Natural Sciences and Engineering Council of Canada (S.E.), the Canada First Research Excellence Fund to BrainsCAN. We also acknowledge the support of the Government of Canada’s New Frontiers in Research Fund (NFRF), [NFRF-T-2022-00051].

## Notes

### Competing Interest Statement

The authors have declared no competing interest.

